# T cell responses induced by attenuated flavivirus vaccination are specific and show limited cross-reactivity with other flavivirus species

**DOI:** 10.1101/2020.01.17.911099

**Authors:** Alba Grifoni, Hannah Voic, Sandeep Kumar Dhanda, Conner K. Kidd, James D Brien, Søren Buus, Anette Stryhn, Anna P Durbin, Stephen Whitehead, Sean A. Diehl, Aruna D. De Silva, Angel Balmaseda, Eva Harris, Daniela Weiskopf, Alessandro Sette

## Abstract

Members of the flavivirus genus share a high level of sequence similarity and often circulate in the same geographical regions. However, whether T cells induced by one viral species cross-react with other related flaviviruses has not been globally addressed. Here, we tested pools of epitopes derived from dengue (DENV), zika (ZIKV), Japanese Encephalitis (JEV), West Nile (WNV), and yellow fever (YFV) viruses by Intracellular Cytokine Staining (ICS) using PBMCs of individuals naturally exposed to DENV or immunized with DENV (TV005) or YF17D vaccines. CD8 T cell responses recognized epitopes from multiple flaviviruses, however, the magnitude of cross-reactive responses was consistently several-fold lower than those to the autologous epitope pools, and associated with lower expression of activation markers such as CD40L, CD69, and CD137. Next, we characterized the antigen sensitivity of short-term T cell lines (TCL) representing twenty-nine different individual epitope/donor combinations. TCL derived from DENV monovalent vaccinees induced CD8 and CD4 T cells that cross-reacted within the DENV serocomplex but were consistently associated with more than 100-fold lower antigen sensitivity for most other flaviviruses, with no cross-recognition of YFV derived peptides. CD8 and CD4 TCL from YF17D vaccinees were associated with very limited cross-reactivity with any other flaviviruses, and in five out of eight cases more than 1000-fold lower antigen sensitivity. Overall, our data suggest limited cross-reactivity for both CD4 and CD8 T cell responses between flaviviruses and has implications for understanding immunity elicited by natural infection, and strategies to develop live attenuated vaccines against flaviviral species.

**Importance:** The envelope (E) protein is the dominant target of neutralizing antibodies for dengue virus (DENV) and yellow fever virus (YFV). Accordingly, several DENV vaccine constructs use the E protein in a live attenuated vaccine format, utilizing a backbone derived from a heterologous flavivirus (such as YF) as a delivery vector. This backbone comprises the non-structural (NS) and capsid (C) antigens which are dominant targets of T cell responses. Here, we demonstrate that cross-reactivity at the level of T cell responses amongst different flaviviruses is very limited, despite high levels of sequence homology. Thus, the use of heterologous flavivirus species as a live attenuated vaccine vector is not likely to generate optimal T cell responses, and might thus impair vaccine performance.

## Introduction

Flavivirus infections can cause a wide variety of clinical manifestations and complications in humans, ranging from undifferentiated fever, vascular leak syndrome, encephalitis and death. Because of their high prevalence worldwide, the four serotypes of dengue virus (DENV), yellow fever virus (YFV), West Nile virus (WNV), Japanese encephalitis virus (JEV), and most recently Zika virus (ZIKV), are responsible for tens of millions of disease cases and thus have a large global impact on human health and disease (4, 21).

Despite intense investigation, the immune correlates of disease and vaccine efficacy are not well defined, particularly in the case of DENV and ZIKV (25). Both antibody and T cell responses have been reported to play a role in immunity and immunopathology (19, 24, 32, 42, 46, 50, 53). Of particular interest in this context are the potential contribution of flavivirus cross-reactive antibodies and T cell responses to both disease protection and immunopathogenesis.

The envelope (E) protein, a major virion surface protein, is involved in receptor binding and membrane fusion and induces neutralizing antibodies in the infected hosts. Human infection results in the production of both virus species-specific and flavivirus cross-reactive antibodies (39). In the case of DENV, most individuals generate cross-reactive antibodies that initially protect against the spread of infection, but may later enhance infection and/or disease with heterologous serotypes (24, 25). Similarly, cross-neutralization in acute ZIKV infection in donors with pre-existing DENV immunity was the strongest in early convalescence but waned to low levels over time (31). JEV vaccination-induced high levels of JEV neutralizing antibodies but also DENV cross-reactive antibodies, which at sub-neutralizing levels, possessed DENV infection-enhancement activity (38).

At the level of T cell reactivity, a similar pattern has been reported, with clear cross-reactivity within different DENV serotypes. It has been proposed that cross-reactive T cells raised against the original infecting serotype dominate during a secondary heterologous infection, a phenomenon that has been termed “original antigenic sin” (20, 29). It was hypothesized that during secondary infection, expansion of pre-existing, lower avidity, and cross-reactive memory T cells may induce a “cytokine storm” contributing to immunopathogenesis (29). In contrast with this initial theory, several lines of evidence suggest that both CD4 and CD8 T cells are involved in resolving DENV infection. It has been demonstrated that both CD4 and CD8 T cells can have a direct role in protection against DENV challenge in a murine model (63, 64) and strong, multifunctional, T cell responses correlated with alleles associated with protection from severe disease in humans naturally exposed to DENV (1, 9, 45, 53, 56). These data implied a protective role for T cells against severe DENV disease (50). Similarly, it has been demonstrated that T cell immunity to ZIKV and DENV induced responses that are cross-reactive with other flaviviruses in both humans (15) and HLA transgenic mice(36).

Vaccines for JEV and YFV, but not for WNV or ZIKV(28) are currently licensed for use in humans, and are based on live attenuated vaccine (LAV) platforms. A DENV LAV, based on a chimeric DENV/YFV, was recently licensed but significant controversy remains over its safety and efficacy (16, 43). While all licensed vaccines rely on serological markers as immune correlates measured with validated assays (25), the potential role of T cell-mediated immunity is not yet fully understood. This is relevant since a general hallmark of LAVs is their ability to induce both humoral and cellular immune memory. We previously defined in the DENV context, the antigens recognized as immunodominant by both CD8 and CD4 responses (11, 50, 51, 54). Non-structural (NS) proteins NS3, NS4B and NS5 were the dominant antigens for CD8 T cell responses, while for CD4 T cell responses, the capsid (C), together with NS2A, NS3, and NS5 were immunodominant (51).

Over the past few years, several full-length live-attenuated vaccines containing antigens from all four DENV serotypes (tetravalent vaccines) have been developed. The National Institute of Allergy and Infectious Diseases has developed the live attenuated dengue vaccines TV003/TV005 containing attenuated DENV1, 3 and DENV4 viruses plus a chimeric DENV2/4 virus (60) while Takeda’s live-attenuated tetravalent dengue vaccine candidate (TAK-3) is comprised of an attenuated DENV-2 strain plus chimeric viruses containing the prM and E genes of DENV-1, -3 and -4 cloned into the attenuated DENV2 backbone (33). Thus, both vaccines would be expected to elicit cellular immunity cross-reactive amongst different serotypes. In fact, T cell responses following tetravalent vaccination with TV005 are focused on the highly conserved NS proteins (49). Likewise, it has been reported that TAK-003, which is based on a DENV-2 NS backbone, induces significant cross-reactive responses against NS proteins of DENV-1, -3 and -4 (48).

The most advanced vaccine against dengue virus, Dengvaxia, was recently licensed, and is based on chimeric viruses containing the prM and E genes of DENV-1, -2, -3 and -4 cloned into the attenuated YF backbone (18). This vaccine has been associated with lower efficacy as well as safety issues (43). In the case of the Dengvaxia vaccine, CD8 cellular immunity will have to rely on YF/DENV T-cell cross-reactivity, since the NS proteins encoded in the vaccine are derived from YFV and not DENV. A potential decreased or compromised cellular immunity might be a potential factor contributing to the lower efficacy. Thus, it is of interest to address to what extent DENV and YFV responses induced by vaccination are cross-reactive.

Here, we characterize immune responses elicited by the TV005 and YF17D vaccines to identify and define the functional attributes of cross-reactive responses at the single epitope level between different flaviviruses. The majority of TV005 induced CD4 and CD8 T cells recognize the DENV serocomplex, while the YF17D vaccine-induced fewer cross-reactive T cells. Characterization of the extent and functionality of CD4 and CD8 T cell cross-reaction across different flaviviruses will contribute to the understanding of immunity in natural infections, and has particular implications for vaccine efficacy and safety in endemic settings.

## Results

### Sequence homology of CD8 epitope pools representative of five prevalent flavivirus species

To address to what extent T cells induced by live attenuated DENV or YF vaccines cross-react with other flaviviruses, we developed pools of several hundred predicted or experimentally defined CD8 epitopes from five prevalent flaviviruses (DENV, ZIKV, YFV, WNV and JEV). The process used to define each of these epitope MegaPools (composed of 9-mers and 10-mers), referred hereafter as MPs, is described in more detail in the materials and methods section. As shown in **Table 1** each MP contained an average of 316 peptides, (ranging from 268-368 peptides/pool) derived from all the ten proteins (C, M/E and NS1-5).

**TABLE 1.**
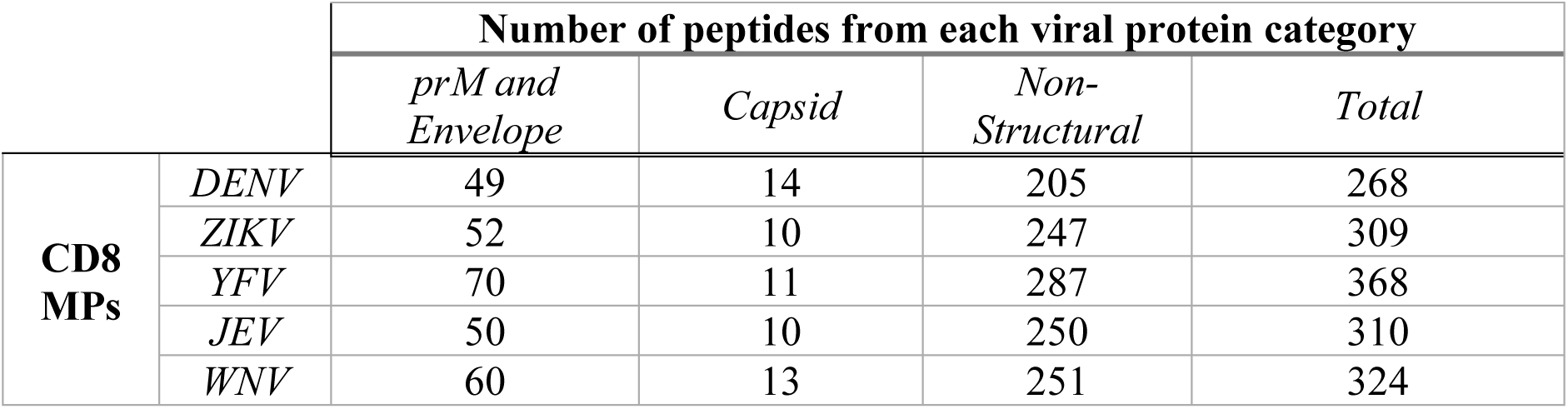
Source Proteins of peptides contained in the flaviviruses MPs.

**Table 2**, lists the number of epitopes for each MP that shared 70% or more sequence identity with DENV, ZIKV, YFV, WNV and JEV consensus sequences, respectively (62). As expected, based on varying degrees of homology between the different viruses, the number of conserved epitopes was highest between DENV and ZIKV, and between WNV and JEV. None of the epitopes included in the various MP shared 70% or more sequence identity with control viral sequences derived from the Ebola virus (EBOV), Chikungunya virus (CHIKV) and Hepatitis C virus (HCV).

**TABLE 2.**
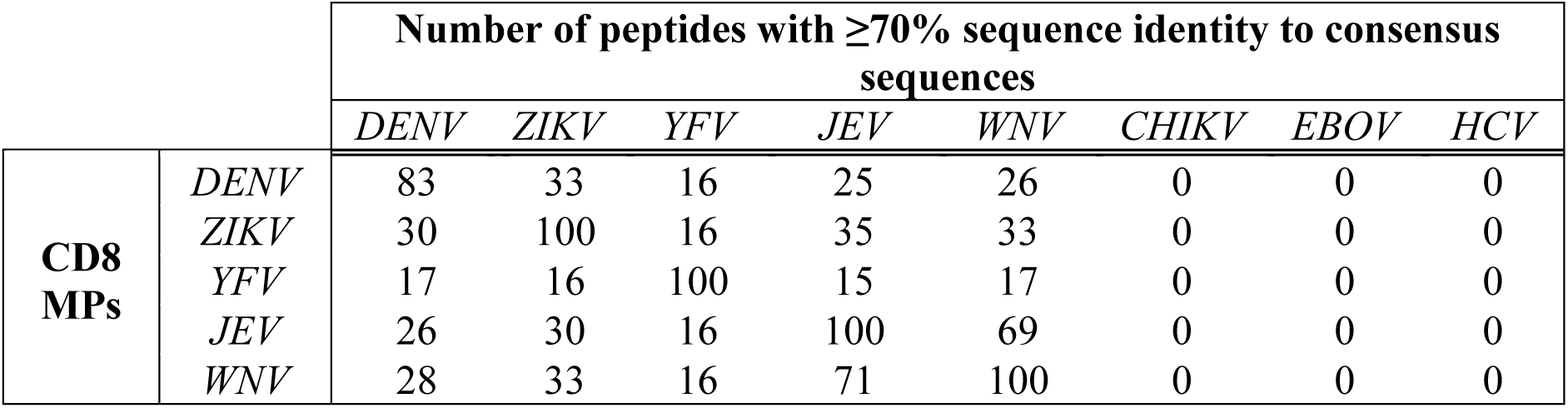
Sequence identity of CD8 flaviviruses MPs and consensus sequences of indicated flaviviruses. The percent of sequence identity in each DENV serotype (DENV1, -2, -3 and -4) was calculated independently and the maximum value was assigned to represent the DENV sequence identity.

| | | Number of peptides with $\geq 70\%$ sequence identity to consensus sequences | | | | | | | |
| --- | --- | --- | --- | --- | --- | --- | --- | --- | --- |
|  |  | *DENV* | *ZIKV* | *YFV* | *JEV* | *WNV* | *CHIKV* | *EBOV* | *HCV* |
| **CD8**<br>**MPs** | *DENV* | 83 | 33 | 16 | 25 | 26 | 0 | 0 | 0 |
|  | *ZIKV* | 30 | 100 | 16 | 35 | 33 | 0 | 0 | 0 |
|  | *YFV* | 17 | 16 | 100 | 15 | 17 | 0 | 0 | 0 |
|  | *JEV* | 26 | 30 | 16 | 100 | 69 | 0 | 0 | 0 |
|  | *WNV* | 28 | 33 | 16 | 71 | 100 | 0 | 0 | 0 |

### Measuring CD8 T cell responses in flavivirus-endemic areas

Addressing the extent of T cell cross-reactivity amongst several flaviviruses is important to understand the potential impact of exposure to multiple subsequent flaviviruses in endemic areas. To address this point, our overall approach was to assess the ability of heterologous flavivirus MPs to elicit the production of IFNγ from memory CD8+ T cell responses in samples from Nicaragua and Sri Lanka. To determine whether DENV specific T cell responses might be cross-reactive with other flavivirus epitopes, we studied peripheral blood mononuclear cell (PBMC) samples from blood bank donors in Managua (n=8) and Colombo (n=6) previously selected to be DENV seropositive and being categorized as high responders against the DENV MP (The high responders were defined by a stimulation Index >2 and background reactivity below 0.1, see Methods). **Fig. 1** shows the reactivity against all five MPs expressed as the percentage of CD3+CD8+ IFNγ-producing cells. As expected due to the selection criteria of those donors, significantly high reactivity was observed after DENV MP stimulation with a geometric mean response of 0.24 (p=0.0001 when compared to the same unstimulated cells as control (CTRL) with paired non-parametric Wilcoxon test). In addition, significant reactivity in DENV MP-reactive donors was observed to ZIKV, YFV, WNV and JEV MPs, with geometric mean in the 0.056 to 0.074 range (p values as compared to control were 0.048 for ZIKV, 0.013 for YFV, 0.037 for WNV and 0.046 for JEV). These results demonstrated that five heterologous MPs recalled a significant response in DENV MP-reactive subjects when compared to an unstimulated control (CTRL).

**FIG 1.**
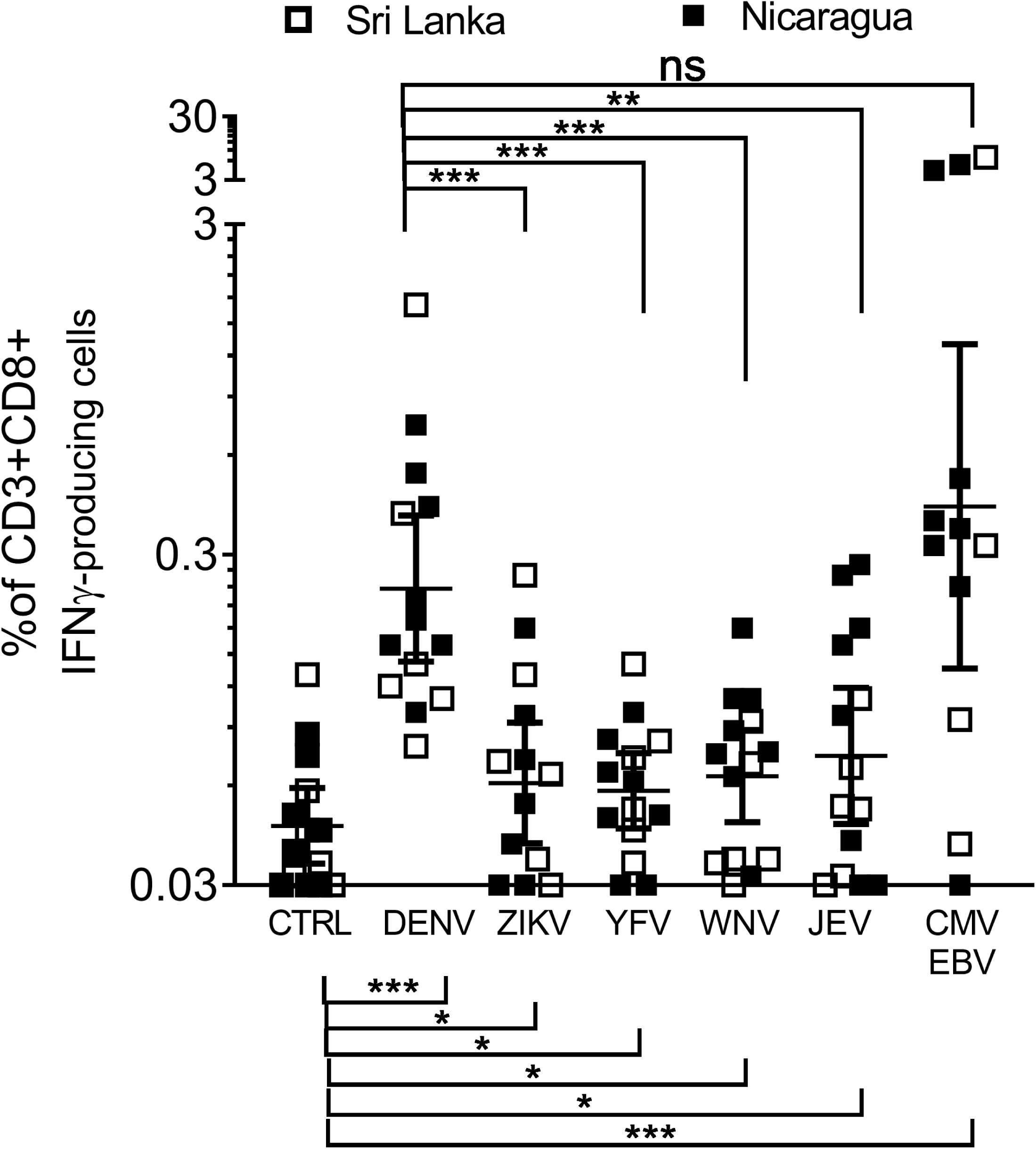
CD8 reactivity against flavivirus MP in flavivirus endemic areas. Percent of CD3+CD8+ IFINγ-producing T cells after flavivirus MP stimulation for 6 hours PBMCs derived from Blood Bank donors in Nicaragua (n=8) and Sri Lanka (n=6). Statistical analyses have been performed using paired non-parametric Wilcoxon test. **p < 0.01, ***p < 0.001, ****p < 0.0001, ns not significant.

To determine the extent of cross-reactivity, we next compared the magnitude of the homologous responses elicited by the DENV MP in DENV-reactive donors with the heterologous responses elicited by the ZIKV, YFV, JEV and WNV MPs and found that the heterologous MP responses were significantly lower in magnitude than the homologous MP responses (p values ranging from 0.0005-0.0479).

To confirm that the results observed were indeed flavivirus-specific, we also assessed T cell reactivity in this cohort against a non-flavivirus CMV/EBV MP. We found a significant T cell response when comparing the CMV/EBV MP to the unstimulated control (geometric mean of the response of 0.42 and p-value of 0.0005) but no significant difference in terms of T cell reactivity when it was compared to the DENV MP (p=0.4697).

These results are compatible with the notion that DENV reactive CD8 T cells responses might recognize certain cross-reactive epitopes contained in the other MPs, although to a significantly lower extent, in terms of magnitude of response as the geometric mean percentage of CD3+CD8+IFNγ+ in each heterologous flavivirus MP is 4 to 5 fold less than the one observed after DENV MP stimulation. As some of these samples were collected in Sri Lanka, an endemic area where other flaviviruses are circulating, it cannot be excluded that some of the response detected was due to exposure to the other flaviviruses. Whether this was indeed the case could not be addressed in the Sri Lanka samples as they were derived from buffy coats from normal blood donations and thus neither clinical history details nor serum samples were available. In Nicaragua, however, these PBMCs were collected before the introduction of ZIKV, and no YFV, WNV or JEV is known to be circulating; additionally, there is no YFV vaccination of the general public. Thus, previous exposure to other flaviviruses is highly unlikely in the Nicaraguan samples.

### Cross-reactivity pattern of CD8 T cell responses induced by a tetravalent dengue vaccine (TV005)

To address potential cross-reactive responses in a controlled exposure setting, and exclude the possibility that previous unknown flavivirus exposure influence the results observed we utilized a cohort of US donors who were vaccinated with experimental tetravalent dengue live attenuated vaccine (TDLAV) candidates TV005, 6-12 months prior to blood collection. All of these donors were confirmed flavivirus naïve before vaccine administration (26).

Specifically, we tested the CD8 T cell IFNγ reactivity against all five flavivirus pools (DENV, ZIKV, YFV, WNV and JEV) in PBMCs derived from TDLAV vaccinated and unvaccinated flavivirus naïve control donors (**Fig 2A**). As expected, no reactivity to either MP was observed in the case of the unvaccinated controls (no significant differences between DENV MP and CTRL groups). Also as expected, the strongest reactivity was detected in the case of TDLAV vaccinees against the DENV MP, with a 0.22 geometric mean response of CD8 T cells producing IFNγ (p=0.0001 compared to the unstimulated control by paired non-parametric Wilcoxon test, and p<0.0001, when compared to the DENV MP-stimulated unvaccinated group with unpaired non-parametric Mann Whitney test).

**FIG 2.**
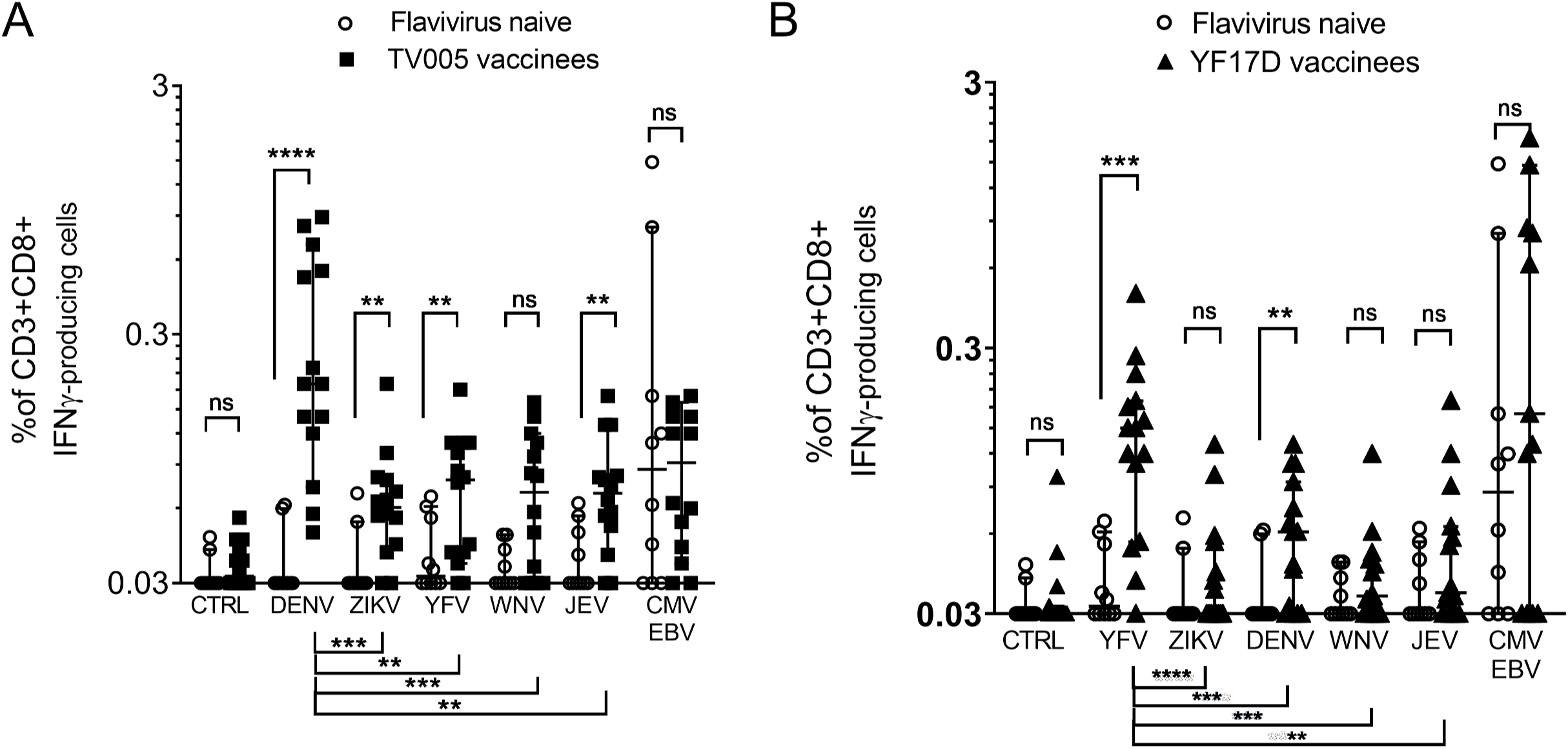
CD8 reactivity against flavivirus MP in vaccination. Percent of CD3+CD8+ IFNγ-producing T cells after flavivirus MP stimulation for 6 hours. A) Reactivity of TV005 vaccinees (n=14) compared to flavivirus naïve (n=10). B) Reactivity of YF17D vaccinees (n= 15) compared to the same flavivirus naïve cohort. Data are expressed as Geometric Mean with 95% CI. Statistical analyses between the different cohorts have been performed using unpaired non-parametric Mann-Whitney test, while statistical analyses for the same cohort across stimuli have been performed using paired non-parametric Wilcoxon test. **p < 0.01, ***p < 0.001, ****p < 0.0001, ns not significant. White circles represent flavivirus naïve, Black squares represent TV005 vaccinees and black triangles represent YF17D vaccinees.

Statistically significant but weak responses in TDLAV vaccinee samples were detected against ZIKV, YFV and JEV MPs, with geometric mean values in the 0.045 to 0.063 range (p values as compared to either the CTRL or the flavivirus-naïve donors were 0.0007 and 0.0088 respectively for ZIKV, 0.0014 and 0.004 for YFV, 0.0002 and 0.005 for JEV) in the case of the TDLAV vaccinees. In the case of the WNV MP, the responses were not significantly higher than the control (p=0.058), and significantly lower than the DENV MP (p=0.016). Finally, the non-flavivirus MP (CMV/EBV) responses were higher than the control, but as expected did not differ between the vaccinated and unvaccinated group (p values 0.0008 and 0.81). Based on these results we conclude that TDLAV vaccination induced CD8 responses that are also capable of recognizing ZIKV, YFV and JEV epitopes.

We next examined the level of cross-reactive responses in terms of magnitude. In all cases, cross-reactive responses in TDLAV vaccinee samples were significantly lower (4.6-fold lower on average compared to DENV MP stimulation; p-values 0.0004, 0.0012, 0.002, and 0.0001 for the ZIKV, YFV JEV and WNV MPs, respectively). Thus, we conclude that TDLAV vaccination induces CD8 responses that are also capable of recognizing ZIKV, YFV and JEV epitopes but to a significantly lower extent (**Fig. 2A**).

### Flavivirus cross-reactive CD8 T cell responses induced by the yellow fever vaccine (17D)

We next asked whether a similar pattern of cross-reactivity might be detectable after vaccination with a different attenuated flavivirus vaccine. Accordingly, we tested the CD8 T IFNγ reactivity against all five flavivirus pools in PBMCs isolated from US donors 6-12 months after vaccination with the live attenuated yellow fever vaccine (YFLAV, YF-17D) and unvaccinated US controls. (**Fig. 2B**).

As expected, little to no reactivity to the YFV MP was observed in the case of the unvaccinated controls (no difference between YFV MP and CTRL groups). By contrast, and also as expected, the strongest reactivity was detected against the YFV MP, with a geometric mean of 0.122 (p<0.0001, when compared to the unstimulated control (CTRL), and p= 0.0002 when compared to the YFV MP-stimulated unvaccinated group).

Responses were noted also in the case of the YFLAV vaccinees (as compared to either the control or the flavivirus-naïve donors) when stimulated with the DENV MP (p value= 0.0003 and 0.0016, respectively). Some responses were also noted in the case of the ZIKV, WNV and JEV MPs, with median values in the 0.027 to 0.049 range. These responses were, in general, significant when compared to CTRL, but not significant when compared to unvaccinated donors (ZIKV: p-values 0.0461 and 0.3446, WNV: p-values 0.0001 and 0.3168, JEV: p-values <0.0001 and 0.1473, respectively). Finally, as previously stated, no significant difference between YFLAV vaccinees and flavivirus-naïve controls were observed in the case of a control MP encompassing epitopes derived from the non-flavivirus CMV/EBV viruses (p-values 0.0046 and 0.3352). Based on these results we conclude that YFLAV vaccination induces CD8 responses that are capable of recognizing DENV epitopes, but only marginally if at all ZIKV, WNV and JEV epitopes.

We next compared the magnitude of the YFV MP responses in YFLAV recipients, with those observed in response to the DENV, ZIKV, WNV and JEV MPs. In all cases, responses against DENV, ZIKV, WNV and JEV MPs were significantly lower (3.4-fold lower on average compared to YFV MP; p-values <0.0001, 0.0006, 0.0001 and 0.0012 for the ZIKV, DENV, WNV and JEV MPs, respectively). We conclude that YFLAV vaccination is able to induce cross-reactive CD8 T cell responses recognizing epitopes derived from other flaviviruses, but the magnitude of such cross-reactive responses is significantly lower compared to homologous YFV-derived epitopes (**Fig. 2B**).

### CD8 T cell cross-recognition of heterologous epitopes is associated with lower expression of activation markers

The results above suggest that heterologous cross-reactive responses are in general weaker when compared to the homologous induced responses when stimulated with peptides variants. We next investigated whether, in addition to the difference in magnitude, we could observe differences in the quality of CD8-specific T cell responses, as represented by activation markers. For this purpose, we analyzed the intracellular expression of CD40L, CD69 and CD137 activation markers in virus-specific CD8 T cells (CD3+CD8+IFNγ+) after stimulation with the various MPs (see **Fig. 8** for gating strategy). The percentage of total CD8 T cells expressing the CD40L, CD69 and CD137 markers (**Fig. 3**, white bars) were compared with the percentage of expression in CD3+CD8+IFNγ+ T cells after DENV CD8 MP stimulation (**Fig. 3**, black bars) or after stimulation with other flavivirus MPs (**Fig. 3**, grey bars).

**FIG 3.**
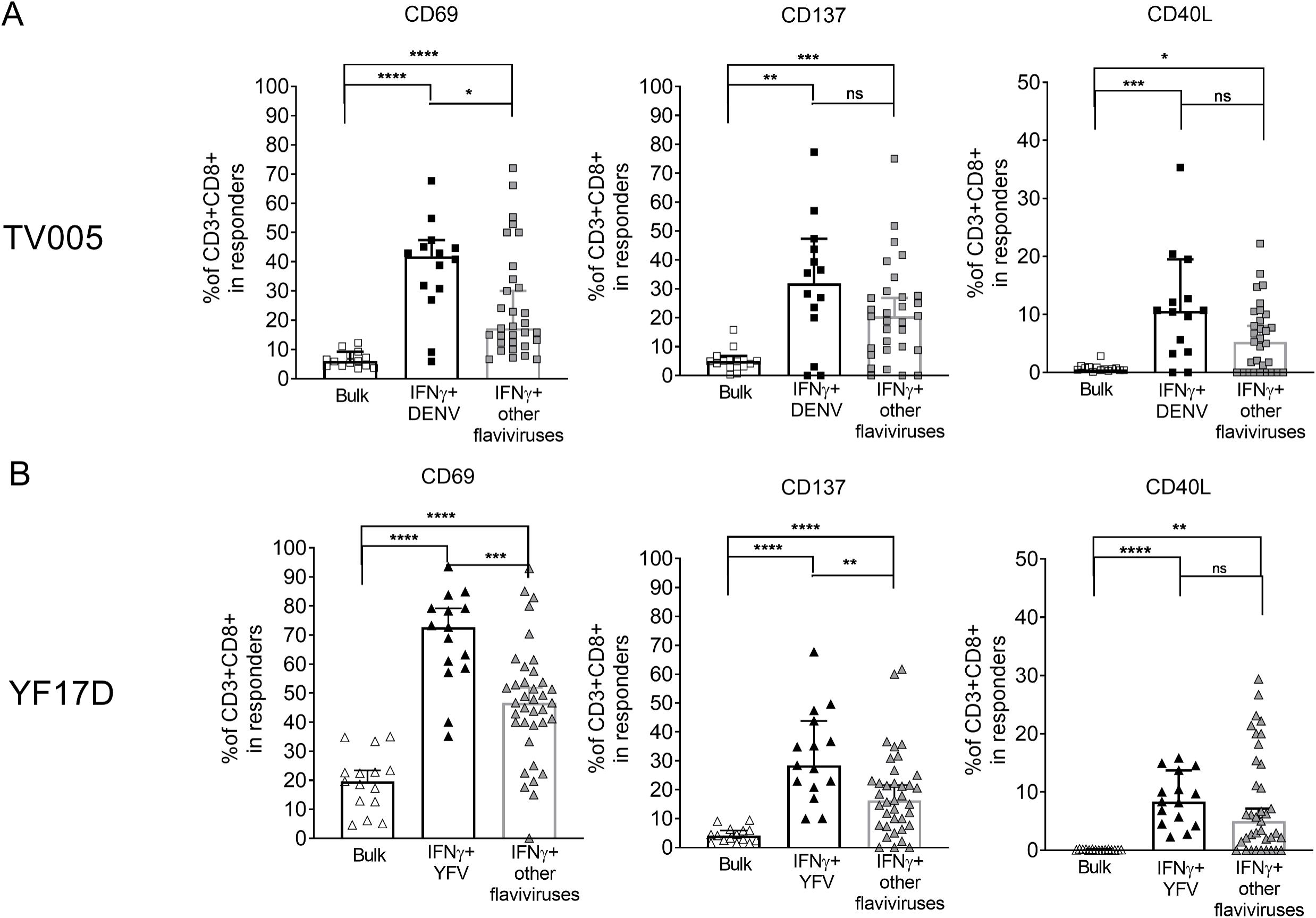
Activation marker expression in IFNγ-producing T CD8 cells after flavivirus specific MP stimulation or cross-reactive flavivirus MP stimulation. Expression of CD69, CD137 and CD40L (for gating strategy see Fig 8) was assessed for individual donor/MP stimulation combinations. Only instances associated with positive responses were examined (defined as % of CD3+CD8+ IFNγ+ above the 0.03 threshold calculated based on mean+2SD of flavivirus naïve MP reactivity). Expression of these markers in the CD3+CD8+ IFNγ+ subset is compared with bulk CD3+CD8+ T cell (white symbols) after stimulation with homologous MP (black symbols) or heterologous MPs (all different heterologous MPs combined; grey symbols). Responses in TV005 vaccinees (A; squares, n=14) or YF17D vaccinees (B; triangles, n=15) are shown. Data are expressed as Median with 95%CI. Statistical analyses have been performed using unpaired non-parametric Mann-Whitney test. *p<0.05, **p < 0.01, ***p < 0.001, ****p < 0.0001, ns not significant.

For TDLAV (TV005) vaccinees, CD69, CD137 and CD40L markers were significantly upregulated in the T cells responding to DENV MP stimulation as compared to the bulk population (CD69: DENV [median = 42%] vs bulk [6%] p<0.0001, CD137 DENV [32%] vs bulk [5%] p=0.0057, CD40L: DENV [11%] vs bulk [0.6%], p=0.0008, Mann-Whitney test). We then compared the expression of these markers on CD3+CD8+IFNγ+ T cells from TDLAV (TV005) vaccinees in response to DENV-specific and heterologous MP stimulation. A significant lower expression of CD69 was detected after stimulation with DENV MP vs other flavivirus MPs, (p= 0.0344) and a non-significant trend was observed for CD40L (DENV-specific vs other flavivirus MPs= 0.0575) and for CD137 (DENV-specific vs other flavivirus MPs= 0.1033) (**Fig. 3A**).

Similarly, YFV-specific homologous stimulation of YFLAV (YF17D) vaccinees was associated with significantly increased expression of CD69, CD137 and CD40L markers as compared to the bulk population (CD69: YFV [73%] vs bulk [0.1%], p<0.0001, CD137: YFV[28%] vs bulk [4%], p<0.0001, CD40L: YFV [8%] vs bulk [0.1%], p<0.0001, Mann-Whitney test; **Fig. 3B**). When we examined the CD3+CD8+IFNγ+ T cell response of YFLAV (YF17D) vaccinees to the other heterologous MPs, we found significantly lower expression for the CD69 and CD137 markers (YFV-specific vs other flavivirus MPs p=0.0006 and 0.0038, respectively), while a non-significant trend was observed for CD40L (YFV-specific vs other flavivirus MPs p= 0.1784) (**Fig. 3B**). Overall, these data suggest that CD8 T cells that recognize cross-reactive heterologous sequences receive less vigorous activation signals, particularly after YFLAV monovalent vaccination.

### Monovalent DLAV vaccination induces CD8 T cell cross-reactivity against other DENV serotypes but is limited against other flaviviruses

The data presented above suggest that DENV or YFV vaccination induces responses that are only marginally cross-reactive with other flavivirus species in terms of both magnitude and activation capacity. These data were obtained utilizing epitope MPs containing hundreds of different peptides. To characterize the phenomenon by a different approach, we analyzed responses against representative individual epitopes. For this purpose, we derived epitope-specific short-term T Cell Lines (TCLs) by stimulating PBMCs for 14 days with the homologous peptide. Their antigen sensitivity was quantified by determining dose-response curves. In parallel, we determined the reactivity of these TCLs to peptides corresponding to the homologous epitope in parallel to their sensitivity to heterologous corresponding sequences derived from the other flaviviruses studied herein. Comparing the dose-response of the homologous epitope with the heterologous peptides from the various flaviviruses allowed for the quantification of relative potencies.

We first determined the level of cross-reactivity in six different TCLs from four monovalent DLAV vaccinees (immunized with either DEN1Δ30 and DEN3Δ30,31). To ensure that the epitopes studied were representative of *in vivo* vaccination, we selected PBMCs and epitopes from donors that we had previously screened in *ex vivo* IFN-γ ELISPOT assays, following vaccination with specific monovalent DLAV vaccines (49). The homologous, as well as heterologous peptides corresponding to the other three DENV serotypes and YFV, ZIKV, JEV and WNV sequences, were tested at six concentrations to assess the relative potency (**Fig. 4 A-F**). As expected in all cases, responses to the homologous peptides were the most dominant. If responses to any of the heterologous peptides were detected, we calculated the fold difference in antigen sensitivity as compared to the homologous peptide.

**FIG 4.**
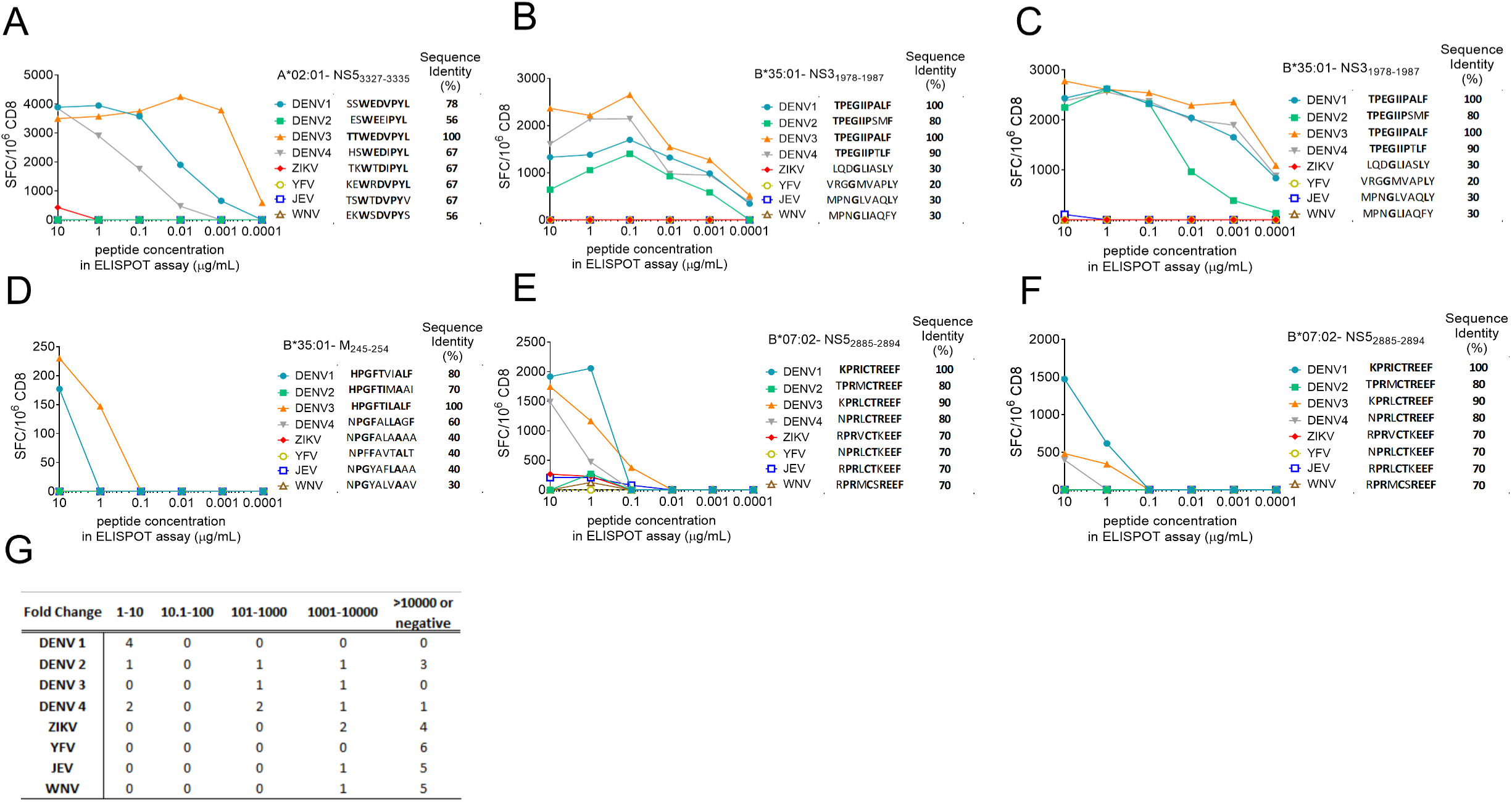
Relative potency of homologous and heterologous flavivirus peptides for CD8+ T cells derived from monovalent DENV vaccination. Spot Forming Cells per million (SFC/10^6) CD8s are plotted for six TCLs stimulated with each peptide at six concentration after 14 days in-vitro expansion and derived from four DENV monovalent vaccinees. Specific peptide responses from vaccinees are shown in A-D (DEN3Δ30,31) and E-F (DEN1Δ30,31); B and C represent independent TCL specific for the same epitopes but derived from two different donors, respectively. G) Summary of the patterns of the relative potency of heterologous peptides compared to the homologous immunizing sequence. Relative potency was calculated for each homologous/heterologous peptide combination based on observed dose responses by recording which peptide dose would give equivalent SFC/10^6^ values. The number of instances where the heterologous sequences were associated with a relative potency of 1-10 (high).10.1-100 (intermediate), >100.1-1000 (weak), 1000.1-10000 (very weak) and >10000 or negative is shown.

Of 42 heterologous peptides tested, high cross-reactivity (defined as reactivity within 10-fold of the homologous peptide) was detected in seven of them (17% of the total). No instance of cross-reactivity was detected in the high and moderate potency range (1-100 fold lower response than to homologous peptide), while in four heterologous peptides (10% of the total) reactivity in the low potency range (101-1000-fold lower response than to homologous peptides) was detected. Finally, seven heterologous peptides (17 % of the total), were associated with very low potency (1001-10000-fold lower response than to homologous peptides) and 24 heterologous peptides (57%) were negative, defined as a more than a 10000-fold lower potency. DENV1 and DENV4 sequences were cross-recognized in the highest number of instances, followed by DENV2. DENV3 showed the weakest relative potency range across all the DENV serotypes. Cross-reactive responses against heterologous ZIKV, JEV and WNV peptides were detected in one peptide each, although with low relative potency (101-1000 fold range), and in all remaining instances, no cross-reactivity was detected. Heterologous YFV peptides did not stimulate cross-reactive T cell responses in all the instances analyzed. From the summary data in **Fig. 4G**, we conclude that, while a degree of cross-reactivity between different DENV serotypes was detected, cross-reactivity with other flaviviruses was limited or, in the case of YFV, totally absent.

### YFV vaccination induces minimal CD8 T cell cross-reactivity against other flaviviruses

To generalize and expand these findings, we performed similar experiments utilizing PBMCs from donors vaccinated with the YF17D vaccine, and epitopes previously identified in *ex vivo* IFN-γ ELISPOT assays, in the context of an epitope identification study (Weiskopf et al, unpublished data). As described above, PBMCs were expanded with YFV-specific epitopes for 14 days. The homologous and heterologous peptides were assayed over a 100,000-fold dose range to assess their relative potency. **Fig. 5A-H** shows results from eight different TCLs derived from six different YF17D vaccinees. For one TCL shown in **Fig. 5C**, cross-reactive responses were detected against all heterologous flavivirus sequences. In this case, the percent of sequence homology between all peptides tested was 90% or more. For the TCLs shown in **Fig. 5B** and **5H**, cross-reactivity with other flaviviruses sequences such as JEV and WNV and DENV2 were detected.

**FIG 5.**
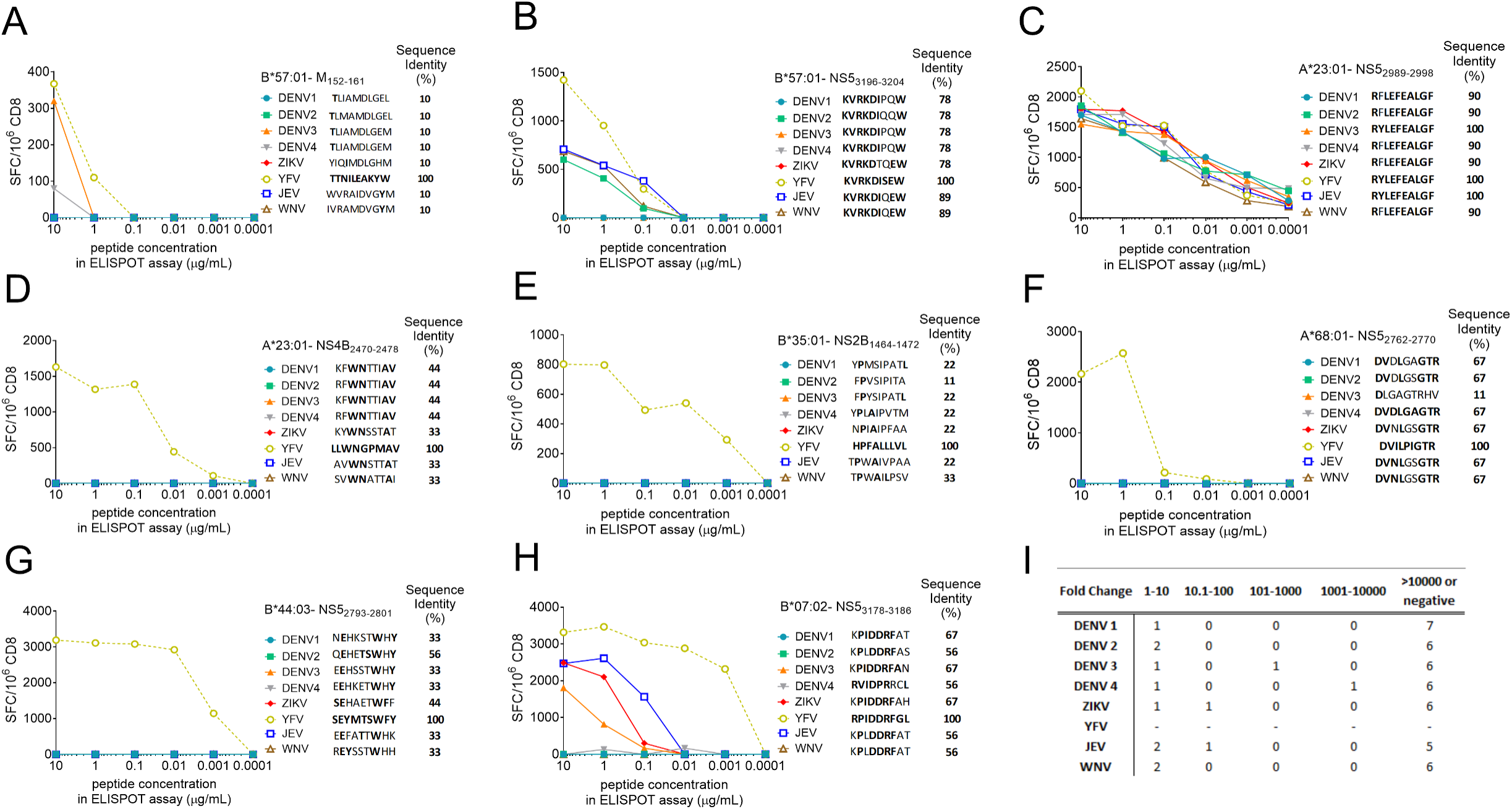
Relative potency of homologous and heterologous flavivirus peptides for CD8+ T cells derived from YF17D vaccination. A-H) Spot Forming Cells per million (SFC/10^6^) CD8s are plotted for TCLs stimulated with each peptide at six concentrations after 14 days in-vitro expansion derived from six YF17D vaccinees. I) Summary of the patterns of the relative potency of heterologous peptides compared to the homologous immunizing sequence. Relative potency was calculated for each homologous/heterologous peptide combination based on observed dose responses by recording which peptide dose would give equivalent SFC/10^6^ values. The number of instances where the heterologous sequences were associated with a relative potency of 1-10 (high).10.1-100 (intermediate), >101-1000 (weak), 1001-10000 (very weak) and >10000 or negative is shown.

Of the 56 heterologous peptides tested, high cross-reactivity (defined as reactivity within a 10-fold of the response induced by the homologous peptide) was detected in ten heterologous peptides (18% of the total). Cross-reactivity in the 10.1-100 fold moderate relative potency range was detected in two heterologous peptides (4% of the total), and only one heterologous peptide was found in the lowest 101 to 1000 fold and 1001-10000 relative potency range (2% of the total in both cases). No cross-reactivity was detected for 42 of the heterologous peptides, corresponding to 75% of the total (**Fig. 5J**). In conclusion, CD8 T cells induced by the YF17D vaccine showed minimal cross-reactivity against other flaviviruses with the DENV serocomplex being the least cross-recognized flavivirus.

### Vaccine-induced CD4 T cell cross-reactivity is even more limited than CD8

Since live attenuated vaccines induce both CD8 and CD4 T cell responses, we asked next whether we could detect cross-reactivity between flaviviruses at the level of CD4 T cell responses. Following the same strategy described above, we expanded DLAV- and YFV-specific CD4 T cell lines for 14 days and tested homologous and heterologous sequences to assess relative potency. We derived six different TCL from two monovalent DENV vaccinees (DEN1Δ30 and DEN3Δ30,31) representing three epitopes recognized in each of the two vaccinees (**Fig. 6A-F**). Of the 45 heterologous peptides tested, high cross-reactivity (defined as reactivity within 10-fold of the homologous peptide) was detected only in two heterologous peptides (4% of the total). More limited cross-reactivity in the moderate 10.1-100 fold lower potency range was detected in four heterologous peptides (9% of the total), and two heterologous peptides in the low 101 to 1000 fold range (4% of the total). Finally, the majority of heterologous peptides (32 out of 45, 71% of the total) were in the very low 1001-10000 lower relative potency range and no cross-reactivity was detected for 5 of the heterologous peptides, corresponding to 11% of the total (**Fig. 6G**). In conclusion, an even lower apparent degree of cross-reactivity between different DENV serotypes was detected within the CD4 compartment compared to the CD8 compartment and the cross-reactivity with other flaviviruses was very limited for both CD4 and CD8 T cells.

**FIG 6.**
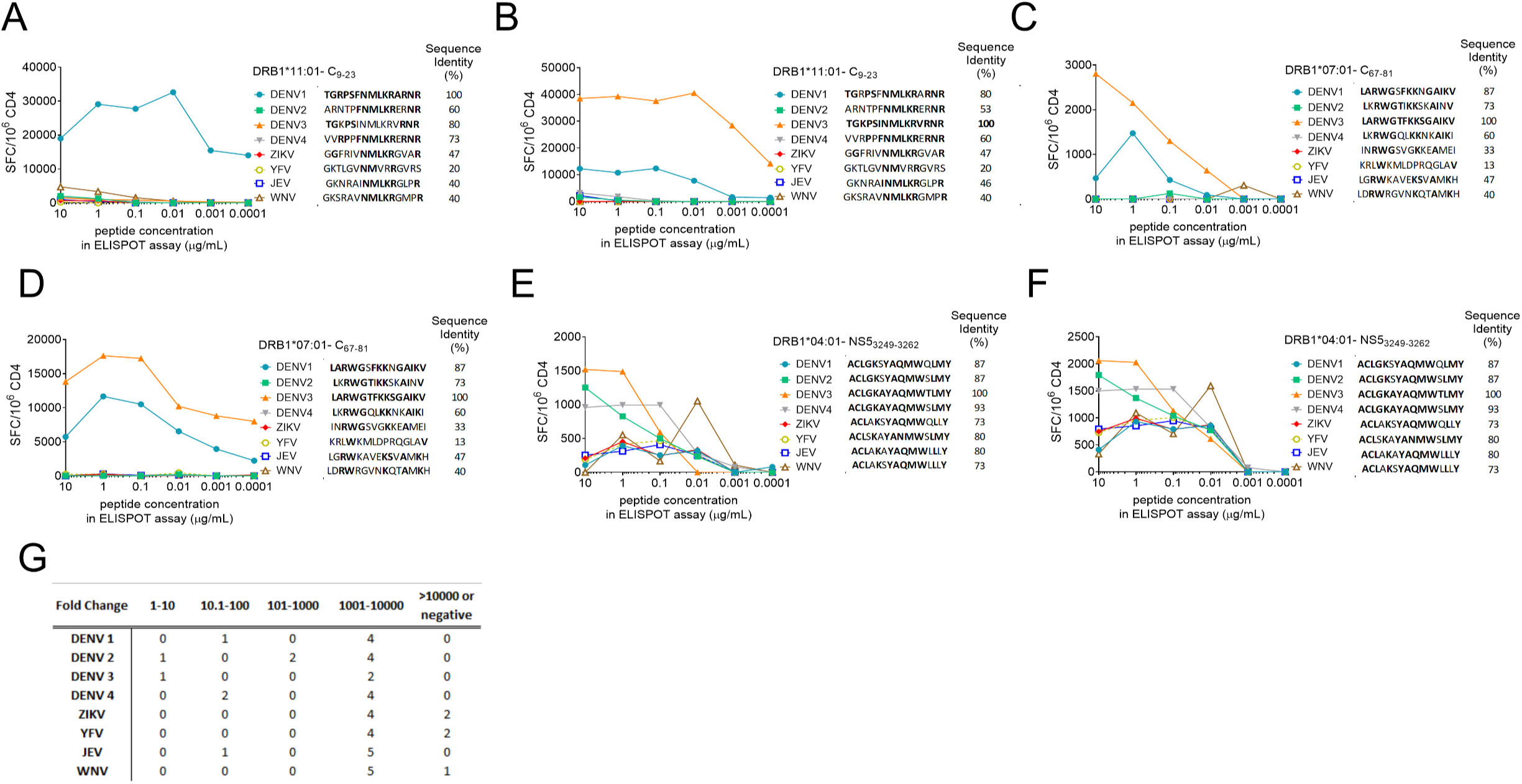
Relative potency of homologous and heterologous flavivirus peptides for CD4+ T cells derived from monovalent DENV vaccination. Spot Forming Cells per million (SFC/10^6) CD4s are plotted for seven TCLs stimulated with each peptide at six concentrations after 14 days in-vitro expansion and derived from six DENV monovalent vaccinees. Specific peptide responses from vaccinees are shown in A (DEN1Δ30,31) and B-F (DEN3Δ30,31); A, B and E, F represent independent TCL specific for the same epitopes but derived from two different donors, respectively. G) Summary of the patterns of the relative potency of heterologous peptides compared to the homologous immunizing sequence. Relative potency was calculated for each homologous/heterologous peptide combination based on observed dose responses by recording which peptide dose would give equivalent SFC/10^6^ values. The number of instances where the heterologous sequences were associated with a relative potency of 1-10 (high).10.1-100 (intermediate), >101-1000 (weak), 1001-10000 (very weak) and >10000 or negative is shown.

Next, we derived TCLs derived from seven different YF17D vaccinees (**Fig. 7A-I**). In seven of the 63 heterologous peptides considered (12% of the total), high cross-reactivity in the 1-10 fold range was observed, all to be ascribed to a single peptide sharing the amino acid core GLYGNG across the different flavivirus species (**Fig. 7E**). Three additional heterologous peptides (0.5% of the total) showed a minimal level of cross-reactivity, all corresponding to the heterologous WNV sequence with relative potency levels within the 1001-10000 range, while the vast majority of the heterologous peptides (82% of the total) did not show any cross-reactivity. Overall, these data demonstrate limited CD4 cross-reactivity against other flaviviruses after YF17D vaccination. In conclusion, our data demonstrate that while vaccination with monovalent DLAV vaccines induced some CD8 and CD4 T cells cross-reactivity, mostly against the other DENV serotypes, the T cell cross-reactivity induced by the YF17D vaccine was limited and mostly absent.

**FIG 7.**
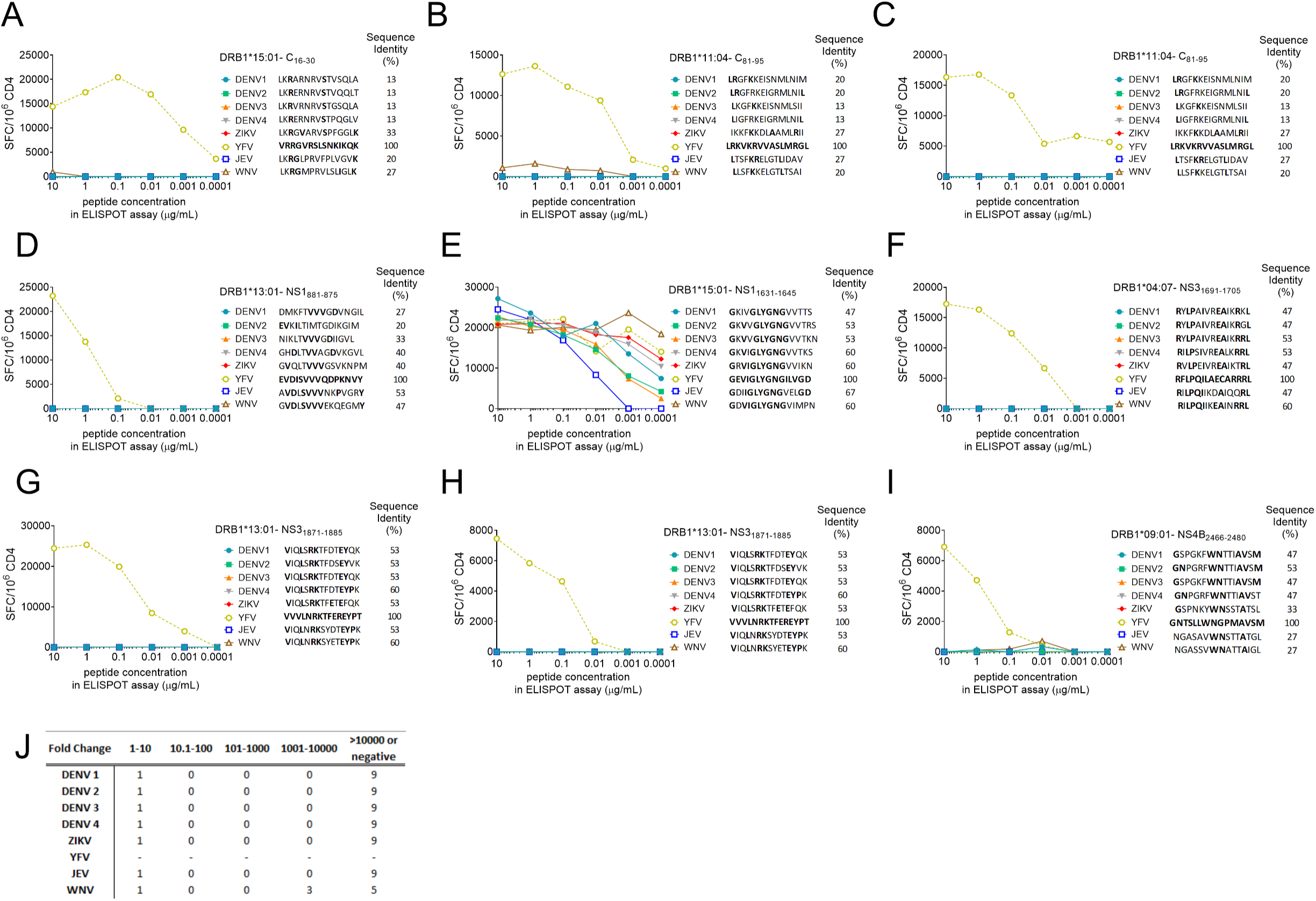
Relative potency of homologous and heterologous flavivirus peptides for CD4+ T cells derived from YF17D vaccination. A-I) Spot Forming Cells per million (SFC/10^6^) CD4s are plotted for TCLs stimulated with each peptide at six concentrations after 14 days in-vitro expansion derived from nine YF17D vaccinees. Specific peptide responses from YF17D vaccinees are shown. J) Summary of the patterns of the relative potency of heterologous peptides compared to the homologous immunizing sequence. Relative potency was calculated for each homologous/heterologous peptide combination based on observed dose responses by recording which peptide dose would give equivalent SFC/10^6^ values. The number of instances where the heterologous sequences were associated with a relative potency of 1-10 (high).10.1-100 (intermediate), >101-1000 (weak), 1001-10000 (very weak) and >10000 or negative is shown.

## Discussion

Flaviviruses such as DENV, ZIKV, JEV, WNV and YFV are highly homologous to each other and often circulate in the same geographical regions. The cross-reactivity is expected to be more pronounced in the case of the TV005 vaccine since in this case it was shown that TV005 focuses the responses on conserved (and thereby by definition cross-reactive) epitopes. Here we studied the level of cross-reactivity of T cells induced by natural infection and vaccination with live attenuated flavivirus vaccines. We demonstrate that broad cross-reactivity amongst sequences of different flaviviruses exists and is largely associated with the recognition of sequences derived from different DENV serotypes. Cross-reactivity amongst different flavivirus species was limited and was associated with responses of lower frequency and magnitude. It was further found that T cells activated from cross-reactive sequences displayed lower levels of expression of common activation markers, and not increased cytokine secretion.

One limitation of this study is that the overall ex-vivo cross-reactive responses were measured after stimulation to selected predicted epitopes MPs. We can not exclude that we missed potential cross-reactive epitopes by mostly including bioinformatically defined epitopes. However, this does not affect the findings at the single epitope level where we tested the cross-reactive potential of experimentally defined epitopes.

These data provide further insights regarding what level of sequence homology is generally associated with potential cross-reactivity.

The data has implications for understanding immunity elicited by vaccination and/or natural infection. In particular, it suggests that the use of YFV as a backbone to engineer live vaccines to deliver E and prM proteins from other flavivirus is not likely to generate optimal T cell responses against other flaviviruses.

The cross-reactivity between DENV- and YFV- derived epitopes observed was fairly limited. The absence of the NS and C DENV proteins, which are immunodominant for CD4 and CD8 responses in Dengvaxia (17), combined with the limited cross-reactivity observed in this study, might contribute to the relatively low level of efficacy observed for this vaccine. This is in agreement with a murine model of heterologous flavivirus infection where previous exposure to YF did not provide cross-reactive functional protection against the DENV1 challenge (39).

Traditionally, flaviviruses have been subdivided into so-called sero-complexes, comprising members that are cross-neutralized by polyclonal sera. This classification largely correlates with the amino acid sequence identity of E and led to the establishment of the DENV sero-complex (consisting of DENV serotypes 1 to 4), and the JEV sero-complex (also including WNV)(22). Zika virus is more closely related to DENV than to the JE virus sero-complex or to YFV which is almost as distantly related to the other mosquito-borne flaviviruses as it is to the tick-borne viruses (22). In accordance with this general overall level of sequence homology, we have observed the highest cross-reactivity within the DENV sero-complex after monovalent DENV vaccination. Thus, while T cell cross-reactivity is appreciable across different sero-complexes/serotypes, T cell cross-reactivity is limited across different flavivirus species.

T cells recognize peptide epitopes derived from the original priming antigen and/or vaccine and are reactivated in subsequent encounters with the same exact epitopes, but also from closely related epitopes. The concept of original antigenic sin, originally described for antibody responses in influenza (23) implies that the pathogen strain shapes subsequent responses to other influenza strains. In the case of DENV, the concept of original sin was postulated to contribute to immunopathology (30, 37), but later studies showed that while previous exposure to different DENV serotypes influenced the repertoire of responding T cells, both in humans (15, 52) and mice (57), the effect was mostly reflected in increases in cross-reactive T cells recognizing conserved epitopes, and cross-reactive T cells were associated with protection in murine models of DENV infection (9, 59). Consistent with this notion, we have previously shown that the simultaneous administration of all four monovalent DENV vaccine strains leads to the induction of highly conserved sequences against all four DENV serotypes (49).

While heterologous sequences were generally associated with incomplete cross-reactivity in this study, this does not rule out a contribution of cross-reactive responses in influencing disease and vaccination outcomes. Indeed, in a murine model of DENV infection it has been shown that despite being associated with lower magnitude responses, cross-reactive CD8 T cell epitopes can still contribute to protection by lowering the viral titer in DENV-infected mice (9, 50, 63, 64). We have also shown that DENV pre-exposure influences T cell responses against the highly homologous ZIKV in both human and murine systems (15, 58).

In our studies, not only the degree of cross-reactive recognition of sequences derived from other flaviviruses was limited, but also the T cell activation induced by the cross-reactive species was suboptimal, resulting in lower expression of several activation markers. This is consistent with the original description by Evavold et al (10) of the phenomenon of Altered Peptide Ligands (APL), epitope variants carrying one or more substitutions. A large body of literature suggests that APL can trigger incomplete T cell activation, and clarified some of the mechanism involves in the effect, as ascribed to lower levels of Zap70 phosphorylation and other TCR signaling alterations (41). Thus, it seems that epitope variants in some cases are fully cross-reactive, while in other cases are incompletely activated. We saw no evidence of increased cytokine production as suggested by other studies hypothesizing a cytokine storm induced by heterologous sequences as a mechanism of DENV pathogenesis (30, 37).

Our data also provides insights regarding what degree of sequence homology is necessary for cross-reactivity, at the level of CD4 and CD8 T cell responses. Specifically, CD8 T cell cross-reactivity was detected in 9 out of 9 instances of heterologous sequences that had one substitution (about 90 % of sequence identity for 9/10-mers) as compared to the immunizing epitope. Cross-reactivity was detected in 6 out of 9 instances of heterologous sequences that had two substitutions (about 80 % of sequence identity for 9/10-mers). Cross-reactivity was detected in 5 out of 15 heterologous sequences that had three substitutions (about 70 % of sequence identity for 9/10 mers). Finally, cross-reactivity was detected in only 5 out of 61 instances of heterologous sequences that had four or more substitution (less than 67 % of sequence identity). Thus, 80 % of cross-reactive responses were associated with 67% or more sequence identity.

This is in agreement with our previous results, where we found that CD8 T cell cross-reactivity was typically detected for heterologous epitopes that shared 70% or higher sequence (52) and substitution of 1-2 amino acids marked the threshold for CD8 epitopes (Weiskopf JI 2011). In contrast, in the case of CD4 responses, no clear pattern could be discerned, with peptides sharing as little as 50% sequence identity being associated with high cross-reactivity. While the molecular mechanism of this difference is not addressed by the current study, this might be related to the fact that in the case of CD4 responses, each antigenic 15mer epitope might bind in several different registers, and as result, the degree of homology of the central core region recognized by the CD4 response might be higher than what is recorded for the overall peptide.

In conclusion, the result of this study emphasizes the need to accurately assess T cell responses and the potential to cross-react with related pathogens in the context of vaccine development and also suggest that when vaccine vectors with significant homology to the vaccine target are used, anti-vaccine vector responses should also be evaluated.

## Material and Methods

### Epitope MegaPool (MP) design and homology analyses

Epitope CD8 MP was produced by sequential lyophilization of flavivirus-specific epitopes as we previously described and in particular, the DENV CD8 MP has been previously generated and validated in DENV exposed individuals derived from different geographical areas (3, 55).

Flavivirus-specific described epitopes were retrieved by querying the Immune Epitope Database (IEDB)(47) utilizing the following search parameters “positive assay only, No B cell assays, No MHC ligand assay, Host: Homo Sapiens and MHC restriction class I”. In the case of ZIKV, YFV, JEV and WNV, experimentally defined epitopes were supplemented by the predicted epitopes using TepiTool (34) algorithm. For this purpose, consensus sequences for Zika (ZIKV), Yellow Fever (YFV), Japanese encephalitis (JEV) and West Nile (WNV) viruses were generated from a multiple sequence alignment of all available strains (taxonomic IDs:64320, 11089, 11072, and 11084) and then blasted to identify the most representative viral isolate per each flavivirus, as we previously described (62).

In the case of YFV, analyses also included the YF17D vaccine strain and a virus isolate deriving from the recent outbreak in Brazil (protein ID: ARM37843.1) (2).

To perform the epitope prediction, a previously described (35) of 27 most frequent A and B alleles was considered, and predictions were performed for both 9-mers and 10-mers with a consensus percentile rank cutoff ≤1.5. A subsequent HLA allele-specific filter was applied based on the percentile cutoff based on our studies performed on DENV infection (50, 55). When the HLA allele considered was not available, the median of the known alleles was used (as summarized in **Table 3**).

**TABLE 3.** List of cutoff used per HLA class I based on previous DENV studies.

| **HLA** | **Percentile**<br>**cutoff** |
| --- | --- |
| *A\*01:01* | 0.75 |
| *A\*02:01* | 0.4 |
| *A\*02:06* | 1.05 |
| *A\*03:01* | 0.35 |
| *A\*23:01* | 1.1 |
| *A\*24:02* | 1 |
| *A\*26:01* | 0.15 |
| *A\*31:01* | 0.85 |
| *A\*33:01* | 0.85 |
| *A\*68:01* | 0.45 |
| *A\*68:02* | 1.5 |
| *B\*07:02* | 0.35 |
| *B\*15:01* | 0.7 |
| *B\*35:01* | 1 |
| *B\*40:01* | 0.25 |
| *B\*44:02* | 0.4 |
| *B\*44:03* | 0.4 |
| *B\*51:01* | 0.6 |
| *B\*53:01* | 0.7 |
| *B\*57:01* | 0.25 |
| *B\*58:01* | 0.3 |
| *Unknown HLA* | 0.6 |

The resulting peptides have been then clustered using the IEDB cluster 2.0 tool and applying the IEDB recommended method (i.e. cluster-break method) with a 70% cut off for sequence identity (5, 6). Peptides were synthesized as crude material (A&A, San Diego, CA), resuspended in DMSO, pooled according to each flavivirus MP composition and finally sequentially lyophilized (3).

Homology analyses to dissect the homology level between each MP and the viral consensus sequences have been performed using the Immunobrowser tool (7). For each MP, the fraction of peptides with a sequence identity of ≥70% with each flavivirus consensus sequence was calculated. In the context of DENV CD8 MP, homology analyses were carried out in each DENV consensus sequences calculated per serotype, then the maximum value of homology obtained across the four serotypes was used for each peptide analyzed.

### Study subjects

PBMCs from DENV endemic areas derived from healthy adult blood bank donors were collected anonymously from National Blood Center, Ministry of Health, Colombo, Sri Lanka, and from the Nicaraguan National Blood Center, Managua, as previously described (12).

In Nicaragua, samples were collected in the 2015-2016 range, prior to the introduction of ZIKV into the Americas. All protocols were approved by the institutional review boards of both LJI and Medical Faculty, the University of California, Berkeley, the Nicaraguan Ministry of Health, and the University of Colombo (serving as NIH approved IRB for Genetech). Blood collection and processing was performed in the two cohorts as we previously described(12, 14).

The yellow fever live-attenuated vaccine (YF17D) cohort and the flavivirus naïve cohorts consist of healthy donors; adult male and non-pregnant female volunteers, 18 to 50 years of age, that were enrolled and either vaccinated with YF17D (n=15) or not (flavivirus naïve cohort; n=10) under the LJI program VD-101.

PBMC deriving from flavivirus naïve and YF17D cohorts were processed in LJI by density-gradient sedimentation using Ficoll-Paque (Lymphoprep; Nycomed Pharma, Oslo, Norway). Isolated PBMC were cryopreserved in heat-inactivated fetal bovine serum (FBS; HyClone Laboratories, Logan UT), containing 10% dimethyl sulfoxide (DMSO) (Gibco) and stored in liquid nitrogen until use in the assays.

The dengue fever live-attenuated vaccinees (TV005) consists of healthy donors, vaccinated with one or four of the dengue live attenuated viruses (DEN1Δ30, DENV4Δ30, DEN3Δ30/31 and DEN2/4Δ30), as previously reported (8, 26, 27, 61). Clinical trials for those vaccinations are described at Clincaltrials.gov under numbers NCT01084291, NCT01073306, NCT00831012, NCT00473135, NCT00920517, NCT00831012, and NCT01072786.

Both vaccinees cohorts were analyzed 6 to 12 months after the initial vaccination.

### Flow Cytometry

Cells were cultured in the presence of either DENV, YFV, ZIKV, JEV, or WNV MPs (1 µg/mL) or DMSO (0.1%) as negative control together with Brefeldin A (BD GolgiPlug, BD Biosciences) for 6 hours. After stimulation, cells were stained with surface markers for 30 min at 4°C followed by fixation with 4% paraformaldehyde (Sigma-Aldrich, St. Louis, MO) at 4°C for 10 min. Intracellular staining was incubated at RT for 30 min after cells permeabilization with saponin, as previously described (13, 44). Detailed information on all the antibodies used for flow cytometry experiments in this study can be found in **Table 4**.

**TABLE 4.** Antibody panel used in flow cytometry experiments to identify both magnitude and quality of CD8+ T Cell response and relevant subpopulations.

| Antibody | Fluorochrome | Volume (mL)/test | Vendor | Catalog | Clone |
| --- | --- | --- | --- | --- | --- |
| CD4 | APC ef 780 | 1 | eBioscience | 47-0049-42 | RPA-T4 |
| CD3 | AF700 | 2 | Biolegend | 317340 | OKT3 |
| CD8 | BV650 | 2 | BioLegend | 301042 | RPA-T8 |
| CD14 | V500 | 2 | BD Biosciences | 561391 | M5E2 |
| CD19 | V500 | 2 | BD Biosciences | 561121 | HIB19 |
| Fixability Dye | ef506 | 1ul/mL of master mix | eBioscience | 65-0866-18 | N/A |
| IFNg | FITC | 1 | eBioscience | 11-7319-82 | 4S.B3 |
| CD154 (CD40L) | PE | 2 | eBioscience | 12-1548-42 | 24-31 |
| CD69 | PE Cy7 | 2 | eBioscience | 25-0699-42 | FN50 |
| CD137(4-1BB) | APC | 2 | BioLegend | 309810 | 4B4-1 |

Surface marker proteins and intracellular cytokine responses were quantified via flow cytometry (LSRII, BD Biosciences) and analyzed using FlowJo software version 10.5.3 (TreeStar Inc., Ashland, OR). The gating strategy is schematically represented in **Fig 8**. Within the CD3+CD8+ subset of lymphocytes, the differences in the magnitude of response between MP stimuli was assessed based on IFNγ+ frequency of parent percentage. The quality of the response was investigated comparing the intracellular staining of CD40L, CD69, CD137 markers within the entire CD8 population and within the CD8+IFNγ+ subsets using background-subtracted values and a 0.03 cut-off for positivity for the different stimuli.

**FIG 8.**
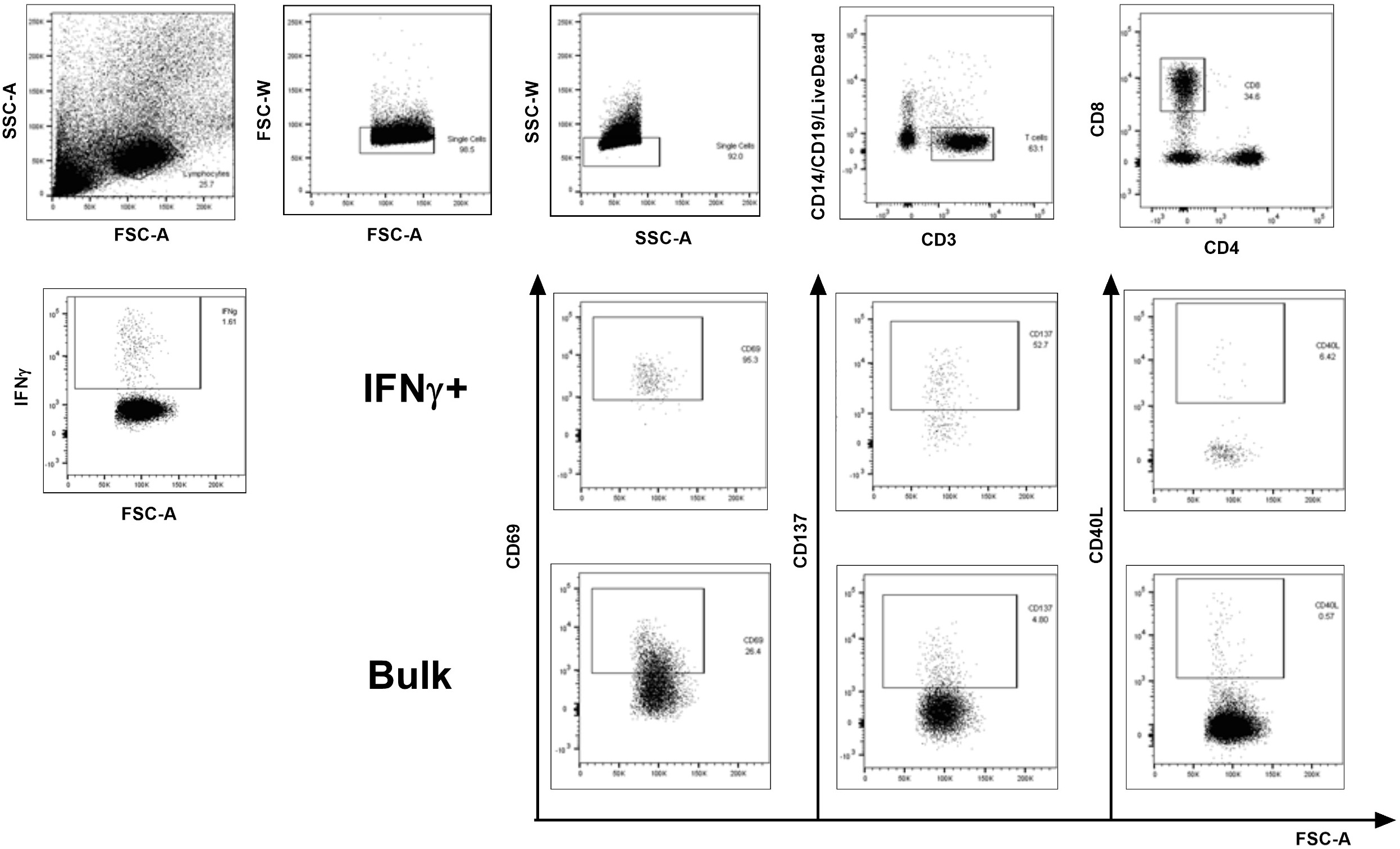
Gating strategy. To examine IFNγ production in CD8+ T Cells, lymphocytes were gated from the whole PBMC on FSC-A and SSC-A axes, followed by exclusion of outlying data points on FSC-W and SSC-W parameters. Cells positive for viability stain, as well as those found to be CD3-were excluded. Of those remaining, cells were separated based on CD4 and CD8 expression parameters and CD8 were exclusively investigated. Intracellular expression levels of T cell activation markers CD69, CD137, and CD40L were examined in whole CD8, “Bulk”, and CD8+IFNγ+ cells. Samples were acquired on an LSRII (BD Biosciences, San Diego, CA).

### ELISPOT assays on short term T cell lines (TCL) to quantitate the antigen dose responses

Short term cell lines for 14 days were set up using donors previously vaccinated with monovalent DENV or YFV vaccines. Cells were expanded using specific DENV epitopes corresponding to the original vaccination and identified in previous studies (49). YFV epitopes were identified using the same approach in YFLAV donors. Cells were expanded using specific DENV/YFV epitopes corresponding to the original vaccination identified using the same approach as previously described (28). After 14 days, IFNγ ELISPOT assays were performed as previously described (12, 50, 51). Briefly, each TCL was tested with the epitope derived from the immunizing vaccine, and peptides corresponding to analogous sequences from the different DENV serotypes or other flaviviruses (YFV, ZIKV, JEV, WNV) in triplicate. Each peptide was tested at six different concentrations (10µg/mL, 1µg/mL, 0.1µg/Ml, 0.01µg/mL, 0.001µg/mL or 0.0001µg/mL). Cells were stimulated for 20hr at 37 C°, 5% CO_2_ at a concentration of 1×10^5^ cells/mL media on plates previously coated with anti-human IFNγ (Mab 1-D1K; Mabtech, Stockholm, Sweden). Cells were then discarded and plates were further incubated with biotinylated IFNγ antibody (Mab 7-B6-1; Mabtech, Stockholm, Sweden) and incubated for 2hr at 37 C°. Avidin Peroxidase Complex (Vectastain ABC Kit, Vector Laboratories, Burlingame, CA) and 3’-amino-9-ethyl carbazole (AEC Tablets, Sigma, St. Louis, MO) were further used to develop the plate. Image analysis was performed using a KS-ELISPOT reader (Zeiss, Munich, Germany).

### Statistics

Statistical analyses were performed using Graph pad Prism (San Diego, CA). Specifically, the analysis of the responses for different cohorts against the same stimuli was performed using unpaired, non-parametric Mann-Whitney test. While to compare the same cohort’s response to different stimuli, a paired, non-parametric, Wilcoxon test was used. The relative potency analyses were performed by determine the dose-response to each homologous peptide required to achieve a level of response that is comparable to the dose-response of the immunizing epitope and calculate the corresponding fold difference in terms of antigen sensitivity determined by measuring the shift in dose-response observed in the x-axis, as previously described (40).

## Acknowledgments

This work was supported by the National Institutes of Health contracts Nr. HHSN272200900042C and HHSN27220140045C to A.S, 75N9301900065 to A.S and D.W as well as P01AI106695 to E.H. Additional support was provided by HHSN272200900045C to SB. The TV003/005 clinical trials were funded by contract HHSN272200900010C of the Intramural Research Program of the NIH, NIAID. AG, HV and CK performed experiments, reviewed data, and planned the experimental strategy. SKD and SB performed and assisted in the bioinformatics analyses. APD, SW, SAD, JDB, EH, AnS and ADDS provided samples and sample information. AG, DW and AS conceived and directed the study, and wrote the manuscript. All authors have critically read and edited the manuscript.

